# Modulation of *Vibrio cholerae* gene expression through conjugative delivery of engineered regulatory small RNAs

**DOI:** 10.1101/2024.03.28.587233

**Authors:** Pilar Menendez-Gil, Diana Veleva, Rita Ramalhete, Brian T. Ho

**Affiliations:** Institute of Structural and Molecular Biology, Department of Biological Sciences, Birkbeck College, London, UK; Institute of Structural and Molecular Biology, Division of Biosciences, University College London, London, UK

**Keywords:** *V. cholerae*, sRNAs, modulation gene expression, T6SS, conjugation

## Abstract

The increase in antibiotic resistance in bacteria has prompted the efforts in developing new alternative strategies for pathogenic bacteria. We explored the feasibility of targeting *Vibrio cholerae* by neutralizing bacterial cellular processes rather than outright killing the pathogen. We investigated the efficacy of delivering engineered regulatory small RNAs (sRNAs) to modulate gene expression through DNA conjugation. As a proof of concept, we engineered several sRNAs targeting the type VI secretion system (T6SS), several of which were able to successfully knockdown the T6SS activity at different degrees. Using the same strategy, we modulated exopolysaccharide (EPS) production and motility. Lastly, we delivered an sRNA targeting T6SS into *V. cholerae* via conjugation and observed a rapid knockdown of the T6SS activity. Coupling conjugation with engineered sRNAs represents a novel way of modulating gene expression in *V. cholerae* opening the door for the development of novel prophylactic and therapeutic applications.

**Importance:** Given the prevalence and spread of antibiotic resistance, there is an increasing need to develop alternative approaches to managing pathogenic bacteria. In this work, we explore the feasibility of modulating the expression of various cellular systems in *Vibrio cholerae* using engineered regulatory sRNAs delivered into cells via DNA conjugation. These sRNAs are based on a regulatory sRNAs found in *V. cholerae* and exploit its native regulatory machinery. By delivering these sRNAs conjugatively along with a real-time marker for DNA transfer, we found that complete knockdown of a targeted cellular system could be achieved within one cell division cycle after sRNA gene delivery. These results indicate that conjugative delivery of engineered regulatory sRNAs are a rapid and robust way of precisely targeting *V. cholerae*.

## INTRODUCTION

The Gram-negative bacterium *Vibrio cholerae* is the causative agent of cholera, a diarrheal disease endemic to several developing countries (1, 2). *V. cholerae* can be found in brackish waters and colonizes human hosts after intake of contaminated water. After colonization of the small intestine, it releases cholera toxin (CT), which is responsible for the disease’s characteristic diarrhoea (3, 4). *V. cholerae* utilizes different mechanisms to ensure survival and colonization of the gut, including the expression of several virulence factors (CT, toxin-coregulated pilus), antibacterial systems (type VI secretion system, T6SS), quorum sensing, and biofilm formation (5).

The current treatment for cholera is the use of rehydration therapies in conjunction with antibiotic treatment (4), however, the emergence and spread of antibiotic resistance in bacteria has prompted an increased effort to develop alternative strategies to combat bacterial diseases. Such efforts include vaccine development (6), inhibitors of quorum sensing (7) or adhesion (8), and use of novel antibacterials like bacteriophages (9) or extracellular contractile injection systems (10). While these approaches can be effective, by seeking to eliminate the pathogen or block their colonization, they create a strong selective pressure for resistant mutants to emerge. An alternative approach likely to be less prone to resistance would be to alter a pathogen’s genetic composition to eliminate its virulence potential rather than the pathogen itself. Indeed, efforts using lysogenic bacteriophages to deliver virulence-neutralizing factors have been successfully demonstrated for *E. coli and Shigella* (11, 12). Unfortunately, the molecular biology and bacteriophage toolbox for *V. cholerae* is not as extensive as for *E. coli*. We therefore wondered if an alternative approach for delivering virulence-neutralizing factors, DNA conjugation, could be used instead. Conjugation systems encoded by broad host-range plasmids have long been used to deliver novel genetic information into *V. cholerae* (13), but there are also *Vibrio*-specific conjugative plasmids that can also be used (14). Additionally, conjugation has been employed as a means of delivering toxic genes for killing pathogens (15, 16).

The virulence-neutralizing factors that have been employed are dCas9 (11) and pathogen-specific transcriptional regulators (12). dCas9 has the disadvantage that it requires the expression of a large dCas9 protein in addition to the guide RNA, while using native transcriptional regulators limits the cellular processes that can be targeted. Instead, we elected to use non-coding regulatory small RNAs (sRNAs) to silence target genes, an approach previously described in *E. coli* (17). sRNAs play a key role in post-transcriptional regulation in bacteria. They act by base pairing to target mRNAs and can regulate mRNA expression both positively and negatively. Most commonly, binding of the sRNA to the target mRNA occurs across the ribosome binding site (RBS) inhibiting ribosome binding and translation, but sRNA binding can also be coupled with degradation by double stranded ribonucleases (18). In *V. cholerae* and other Gram-negative bacteria, RNA chaperone Hfq, will help stabilize the sRNA and help it bind its target (19–21), though not all regulatory sRNAs require Hfq for their function (22).

In this study, we have engineered sRNAs based on the sRNA TarB (22) to modulate *V. cholerae* gene expression. Several sRNAs targeting type VI secretion system (T6SS) activity were able to successfully knockdown T6SS activity to different degrees with the most effective one knocking down T6SS activity nearly to the level of a gene knockout mutant. We also observed that modifying the recognition sequence in the mRNA did not significantly affect the efficiency of the sRNA knockdown effect so long as the overall secondary structure of the sRNA was preserved. We also employed this sRNA silencing strategy to target other processes in *V. cholerae,* including modulating exopolysaccharide (EPS) production and motility. Lastly, we successfully delivered an sRNA targeting T6SS into *V. cholerae* via conjugation and observed a rapid knockdown of the T6SS activity. Coupling conjugation with engineered sRNAs represents a novel way of modulating gene expression in *V. cholerae* opening the door for the development of novel prophylactic and therapeutic applications.

## MATERIAL AND METHODS

### Strains, plasmids, and growth conditions

The strains and plasmids used in this study are listed in Tables S1, and S2 respectively. *Vibrio cholerae* and *Escherichia coli* strains were grown in Luria Bertani broth (LB, 10 g/L tryptone, 5 g/L yeast extract (Formedium), 5 g/L sodium chloride (Fischer Scientific)) at 37°C shaking. When necessary, the following antibiotics were added: streptomycin (50 µg/mL) (Apollo Scientific), chloramphenicol (*V. cholerae* 3 µg/mL, *E. coli* 15 µg/mL) (Sigma-Aldrich) and/or gentamicin (10 µg/mL) (Sigma-Aldrich). *E. coli* MFDpir strains were supplemented with 0.28 mM DAP (diaminopimelic acid) (Sigma-Aldrich). When indicated, cultures were supplemented with 0.2% glucose (Fischer Scientific)) or 0.2% arabinose (Sigma-Aldrich).

### sRNA design

The sRNA sequences and all relevant characteristics can be found in File S1 and Table S3, respectively. Engineered sRNAs were constructed using TarB sRNA as a scaffold (22). The recognition binding sequence was changed to base pair to the specific target genes. This sequence was further modified to maintain (as much as possible) the secondary structure of the original TarB sequence. Vienna RNAfold web server (23) was used to predict sRNA secondary structures and VARNA applet for drawing the RNA structure (24). To confirm the sRNA-mRNA interactions we used the INTaRNAv2 software (25). sRNAs were then synthesized with PCRs using overlapping primers and then cloned into pBAD33 with specific restriction enzymes.

### Plasmid construction

Plasmids were constructed by amplifying inserts by PCR using specific oligonucleotides and the Q5 high fidelity enzyme (New England Biolabs). PCRs were purified using NucleoSpin® Gel and PCR Clean-up kit (Macherey-Nagel). Plasmids and PCRs were digested with the appropriate restriction enzymes (New England Biolabs), purified, and ligated using Instant Ligase Sticky-end Ligase Mix (New England Biolabs). Ligations were transformed into NEB10β competent cells following the manufacturer recommendations (New England Biolabs). Correct plasmids were verified by PCR and sanger sequencing. They were then introduced into *E. coli* donor strains by electroporation and then into *V. cholerae* strains through conjugation.

### Competition Assays

Prey and predators were grown overnight (ON) at 37°C shaking in LB with the appropriate antibiotics. Next day, 1mL of each culture was centrifuged at 5000g for 3 min and pellets were resuspended in 1 mL of fresh LB. 100 µL of each sample were subcultured into 10 mL of fresh LB and were grown for 2h 30 min. The subcultures of strains carrying pBAD33 plasmids were grown with 0.2% arabinose (Sigma-Aldrich). OD_600_ was measured, cultures were centrifuged at 5000g for 3 min and cell density was adjusted to OD_600_=10. Prey and predators were mixed to a ratio of 1:1 and 5 µL of the mixture was spotted twice (technical replicates) in LB +0.2% arabinose plates. Plates were incubated at 37°C for 2h. Then, bacterial spots were cut and resuspended in 1 mL LB. Bacterial suspensions were serially diluted up to 10^-6^ and 5 µL of each dilution was spotted onto selective plates. Plates were incubated at 37°C ON. Next day, colonies were counted, and photos were taken of the plates. At least 3 biological replicates, each in technical replicates, were performed.

### Motility assays

Strains were grown in LB ON at 37°C shaking with the appropriate antibiotics. Next day, OD_600_ was measured, cultures were centrifuged at 5000g for 3 min and cell density was adjusted to OD_600_=1. 1 µL of each sample was spotted in a tryptone (1% tryptone (Fischer Scientific), 0.5% NaCl (Fischer Scientific)), soft 0.3% agar (Formedium) plate containing 0.2% arabinose and 3 µg/mL chloramphenicol. Plates were incubated at 28 °C for 20-22 hours. Next day, halo diameters were measured, and photos were taken. As a control, all samples were serially diluted, spotted into Streptomycin plates, and incubated at 37°C ON. Next day, colonies were counted to ensure equally amounts of all samples. Each experiment was conducted in technical duplicates and repeated at least 4 times.

### Visualization of T6SS activity with fluorescence microscopy

Strains were grown in LB ON at 37°C shaking with the appropriate antibiotics. Next day, 20 µL of each ON culture were subcultured into 2 tubes containing 2 mL fresh LB, one containing 0.2% glucose and the other 0.2% arabinose. After 2h 30 min at 37°C, 1 µL of each culture was spotted into a microscope slide containing a 1.5% agarose (Fischer Scientific) pad made from PBS (Phosphate buffered saline). Once the culture spot was fully absorbed into the agar, 2-minute-long time-lapse recordings were made with image acquisition every 20s. Two to four biological replicates were performed for each experiment. Foci were counted using “Spot counter” while “Find maxima” was used to count cells, both from Image J. Quantification was done in at least 6 different time-lapse videos from at least two biological replicates.

### Visualization of sRNA delivery using fluorescence microscopy

Donor and recipient cells were grown in LB ON at 37°C shaking with the appropriate antibiotics. Next day 20 µL were subcultured into 2 mL of fresh LB and were grown for 2h 30 min. OD_600_ was measured, cultures were centrifuged at 4500g for 10 min and cell density was adjusted to OD_600_=1.5. Donor and recipient cells were mixed to a ratio of 10:1 and 2 µL of the mating was spot into a microscope slide containing an M9 (M9 salts (Sigma-Aldrich), 500M mM MgSO_4_, 40 mM CaCl_2,_ 0.4% casamino acids (Formedium)) +DAP agarose (1.5%) pad. The slide was incubated at 37°C for 1h and 30 min before visualizing the cells under the microscope, as described in the above section. Three independent replicates were conducted and at least 50 conjugation events per biological replicate were analysed.

### Microscopy imaging and analysis

Microscopy was done using a Nikon ECLIPSE Ti2 inverted microscope with CoolED pE4000 illuminator and a Zyla 4.2 Megapixel Camera. Images were recorded using Nikon Elements software and analysed using the temporal-color code plugin of the Fiji software package (26).

## RESULTS

### Engineered sRNA TarVipA shutdowns *V. cholerae* T6SS

*V. cholerae* employs several non-coding sRNAs as part of its virulence regulation program (20). One of these sRNAs, TarB, is encoded in the *Vibrio* Pathogenicity Island and directly regulates the secreted colonization factor TcpF by binding to the 5’ UTR of the *tcpF* transcript in an Hfq-independent manner (22). The TarB binding target includes the ribosome binding site (RBS) (Figure 1A), and likely works by blocking TcpF translation. Using the TarB sRNA as a scaffold, we designed novel regulatory sRNAs targeting different biological processes in V. *cholerae*. As an initial proof of concept, we decided to target the T6SS because of the clear and robust reporters for T6SS activity (27, 28). We maintained the overall secondary structure and the Rho-independent terminator but replaced the target recognition sequence (Figure 1B). Instead of targeting *tcpF*, this new sequence targeted the equivalent region, including the positioning of the RBS and start codon, of the first gene of the major T6SS gene cluster, *vipA*. We named this sRNA TarVipA.

**Figure 1.**
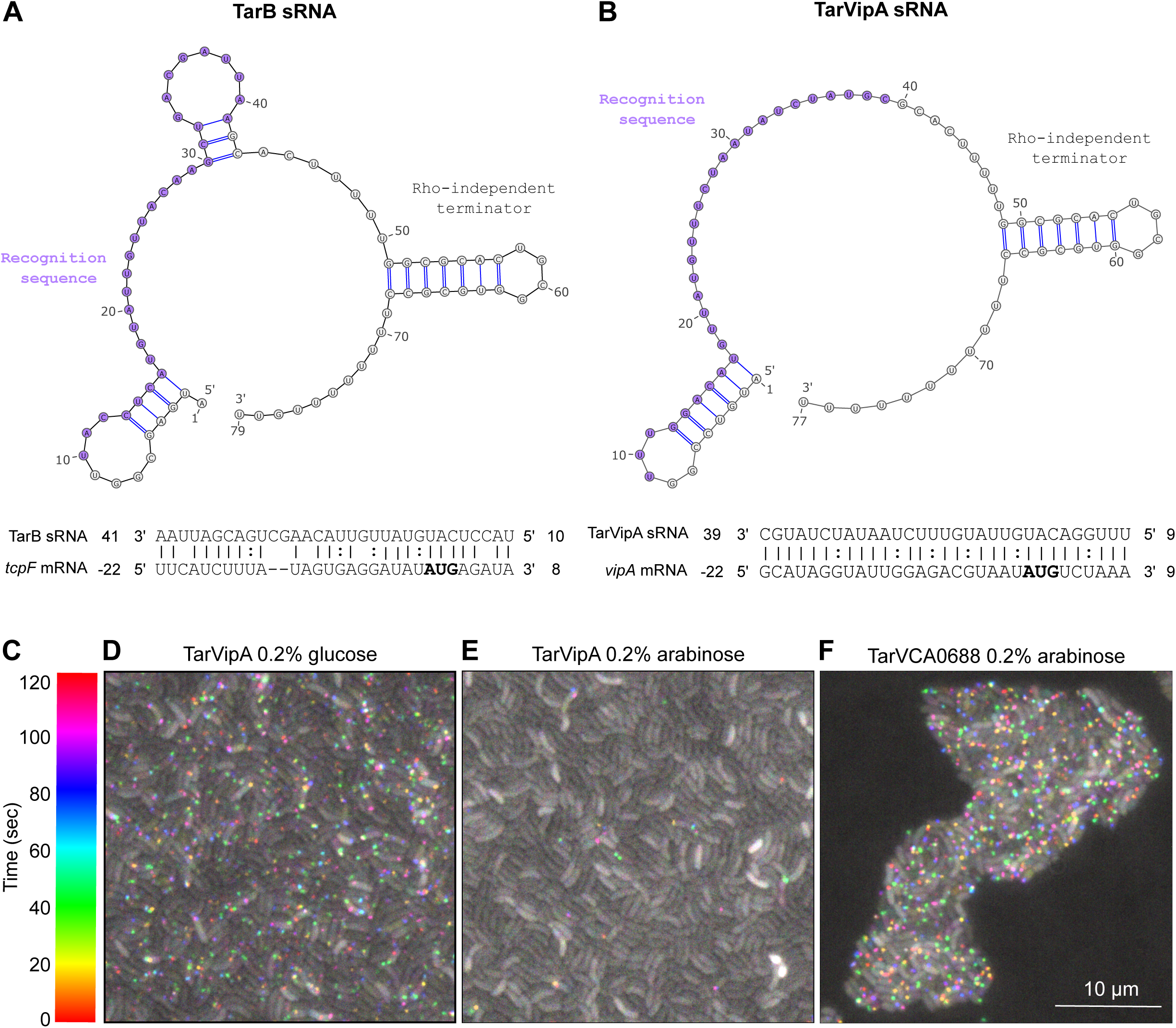
TarVipA sRNA shuts down the T6SS activity of *V. cholerae*. **A)** TarB secondary structure predicted with Vienna RNAfold web server (23) and TarB and *tcpF* mRNA predicted interaction as stated by Bradley et. al. (22). **B)** TarVipA secondary structure predicted with Vienna RNAfold web server (23) and INTaRNAv2 (25) predicted interaction of TarVipA and *vipA* mRNA. The recognition sequences of TarB to *tcpF* mRNA and TarVipA to *vipA* mRNA are highlighted in purple. *tcpF* and *vipA* start codons are highlighted in bold. The numbering of *tcpF* and *vipA* mRNAs is relative to the start of translation whereas TarB and TarVipA numbering is relative to the start of transcription. **C)** Spectrum temporal-colored code from Image J-Fiji software (26) used to analyse the microscopy in D-F. The temporal code assigns a different color to foci appearing in each time point of the time-lapse. White is assigned to non-dynamic foci/aggregates. **D-F)** Time-lapse fluorescence microscopy imagining of *V. cholerae 2740-80 clpV::clpV-mCherry* carrying pBAD33-TarVipA **(D-E)** or pBAD33-TarVCA0688 **(F)**. Cells were either grown in glucose **(D)** or in arabinose **(E-F)** to repress or express pBAD33 expression, respectively. Images were taken every 20s for 2 min. Scale bar is 10 μm.

TarVipA was cloned into plasmid pBAD33 under the control of an arabinose inducible promoter and introduced into *V. cholerae 2740-80* expressing a ClpV-mCherry fusion (28). ClpV-mCherry forms dynamic fluorescent foci as ClpV coalesces onto contracted T6SS sheath structures to facilitate the recycling of secretion apparatus components. In time-lapse, these foci manifest as “blinking” spots and can be used as an indicator of T6SS activity (29). Using a temporal color code (Figure 1C), time-lapse recordings can be projected into a single image enabling the differentiation of dynamic foci (colored spots) from non-dynamic fluorescent aggregates (white spots).

When expression of TarVipA was suppressed (0.2% glucose), colored spots indicating active T6SSs could be observed in nearly every cell (Figure 1D). However, when the TarVipA was expressed (0.2% arabinose), T6SS activity was almost completely eliminated from the population (Figure 1E). As a control we also tested sRNAs targeting non-T6SS related genes and observed wild type levels of T6SS activity for all of them. sRNAs targeting unrelated genes resulted in no elimination of T6SS activity. A representative example, TarVCA0688, targeting the *VCA0688* gene, is shown in Figure 1F.

### Significant positional variation in the specific sRNA target site is tolerated

We next wondered how much flexibility there was for designing the specific TarVipA recognition sequence in terms of shifting the recognition sequence relative to the RBS. To address this, we designed six TarVipA variants in which we shifted the recognition sequence 1, 2, or 4 nucleotides upstream (TarVipA_P+1, TarVipA_P+2 and TarVipA_P+4) or downstream (TarVipA_P-1, TarVipA_P-2 and TarVipA_P-4) relative to the original TarVipA target sequence (Figure S1A). These sRNAs were each cloned into the pBAD33 and introduced into *V. cholerae* 2740-80. Competition assays were performed for each of these *V. cholerae* strains against *E. coli* MG1655 (Figure S1B, C). Expression of the original TarVipA sRNA and five of the six shifted target sRNAs resulted in almost complete knockdown of T6SS activity. However, there was roughly 1-log less survival of the *E. coli* prey compared to a T6SS genetic deletion consistent with our observation of residual T6SS activity in some cells (Figure 1E).

One of our shifted target sRNAs, TarVipA_P+2, was unable to knockdown T6SS activity. After closer examination of the predicted secondary structure of this sRNA, we noticed that its recognition sequence was probably sequestered in a larger hairpin structure (Figure S2). This secondary structure likely prevents TarVipA_P+2 from binding to *vipA* transcript. Overall, these results indicate that sRNA design can tolerate significant shifts in the precise target sequence location, so long as the original secondary structure is preserved.

### The effectiveness of the sRNAs differs depending on the target gene

Having successfully knocked down T6SS activity through targeting of *vipA*, we next looked to see if we could similarly knock down T6SS activity by targeting other essential genes both in the main gene cluster as well as in the auxiliary gene clusters (30, 31) (Figure 2A). Notably, we wondered if our ability to target *vipA* was dependent on it being the first gene in the operon. To this end, we designed sRNAs targeting *vipB*, *tssG*, *vasH*, *tssM*, *hcp*, and *vgrG-2* (Table S3, File S1), all of which have been previously shown to be essential for T6SS activity (27). Each sRNA was cloned into pBAD33, introduced in *V. cholerae*, and their impact on T6SS activity was measured using competition experiments with *E. coli*. Figure 2B shows a representative image of the *E. coli* MG1655 CFUs (colony forming units) recovered (quantified in Figure 2C). Although targeting *vipB*, *tssG*, and *hcp* resulted in a significant knockdown of T6SS activity relative to wild type T6SS, none were as effective as the TarVipA sRNA. Additionally, targeting *vasH*, *tssM*, and *vgrG-2* was no more effective than the control TarVCA0688 sRNA (Figure 2B-C). It is not entirely clear why these disparities were observed but considering that the sRNAs were expressed for at least 6-7 doubling times, differences in protein turnover rate should not be a factor. More likely, the positioning of the gene within the operon and access to target site on the mRNA transcript are going to be the primary factors.

**Figure 2.**
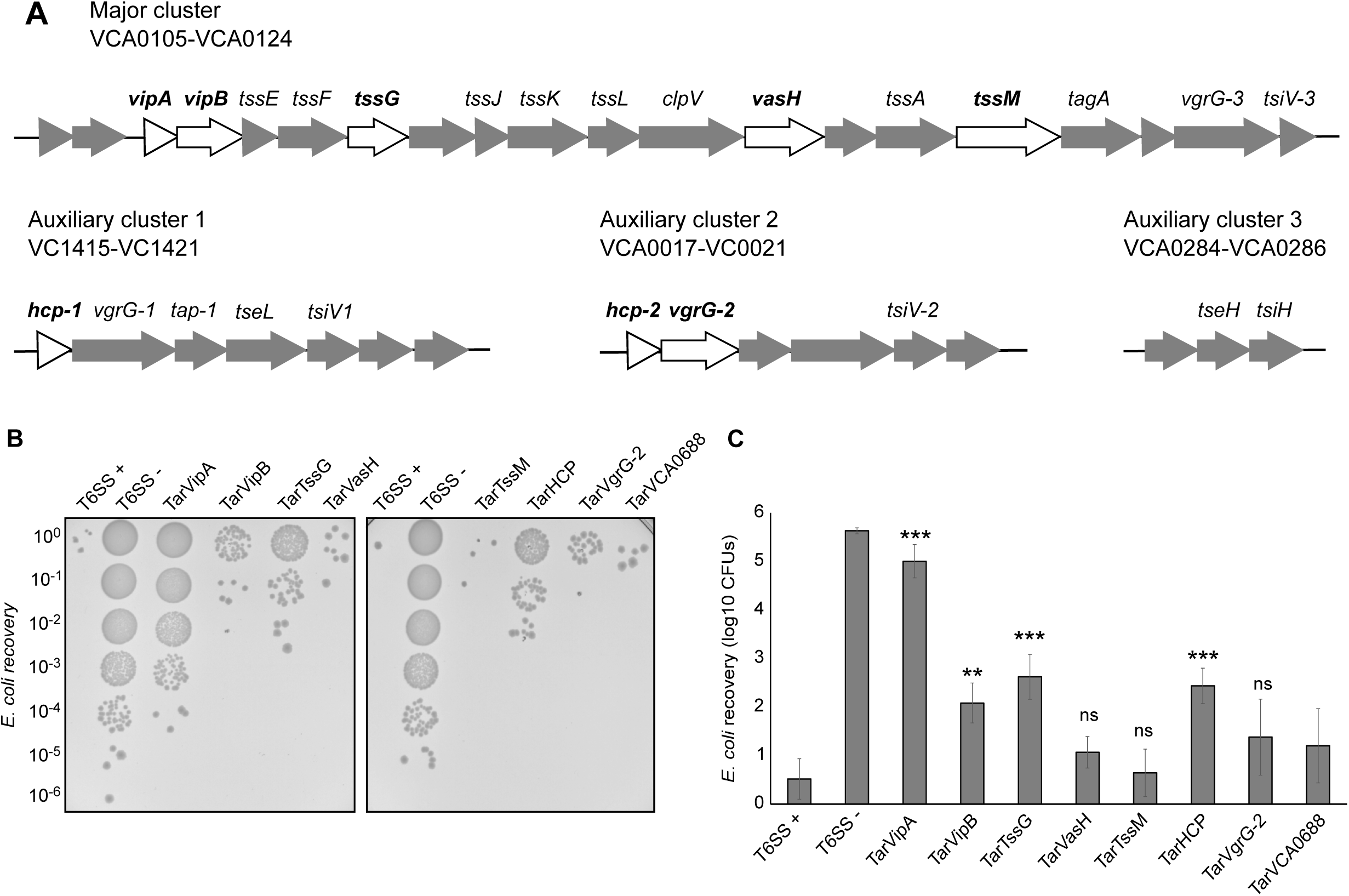
T6SS activity modulation differs depending on the gene being targeted. **(A)** Genetic organization of the T6SS genes. Figure adapted from Joshi et. al. (42). Genes represented in white were targeted by the engineered sRNAs. **(B)** Representative image of the competition assay between *E. coli* MG1655 (prey) and *V. cholerae* 2740-80 *clpV*::*clpV*-*mCherry* strains carrying the sRNAs (predators). As a negative control the predator *V. cholerae* 2740-80 *clpV*::*clpV*-*mCherry* (T6SS +) was included. As a positive control V. cholerae *2740-80 clpV*::*clpV*-*mCherry ΔvipA* (T6SS -) was used. **(C)** Quantification of the *E. coli* CFU recovery after competition with *V. cholerae* strains shown in panel B. Data represent the average of at least three independent replicates, each one done in technical duplicates. Error bars represent the standard deviation (SD) of these three replicates. Asterisks represent statistical significance when compared to the control sRNA (TarVCA0688) sample (paired two-tail t-test, ** p value <0.01, ***p value <0.001; ns not significant).

### Engineered sRNAs can be used to modulate diverse biological processes in *V. cholerae*

Next, we wondered if other cellular processes could be modulated by similarly engineered sRNAs. Exopolysaccharide (EPS) is one of the main components of biofilm in *V. cholerae* and plays a major role in transmission and intestinal colonization as well as survival in aquatic environments (32). EPS is synthesized by several proteins encoded in two regions: *vps-I* (coding for *vpsU* and *vpsA-K*) and *vps-II* (coding for *vpsL-Q*) (32, 33). We designed sRNAs targeting three different genes previously shown to be essential for EPS production: *vpsU* (TarVpsU)*, vpsA* (TarVpsA), and *vpsL* (TarVpsL) (Table S3, File S1). *vpsU* and *vpsA* are the first two genes of the *vps-I* cluster while *vpsL* is the first gene of the *vps-II* cluster (32, 33).

EPS formation was previously shown to protect *V. cholerae* from exogenous T6SS attack (34). We used this property to test the efficacy of sRNAs knockdown. However, when we transformed each of these sRNAs into *V. cholerae V52*, the prey strain used in that study, we did not observe any effect on T6SS sensitivity (Figure 3A). Recognizing that *V. cholerae V52* is an O37 serotype strain with a distinct evolutionary lineage from the *El tor* O1 strains in which TarB has been previously studied, we also introduced each of the sRNAs into a T6SS effector immunity protein knockout mutant of *V. cholerae* C6706. When this strain was mixed with T6SS-active *V. cholerae* 2740-80, two of the sRNAs had a profound impact on their resistance to T6SS-mediated killing (Figure 3A). Although targeting *vpsL* had no effect, TarVpsU expression significantly increased the sensitivity of C6706 to being killed, indicating successful EPS production knockdown. Unexpectedly, bacteria carrying TarVpsA actually became more resistant to T6SS attack, suggesting that engineered sRNAs could also positively regulate EPS production. As a final test, we also introduced the sRNAs into Classical biotype O1 strain, O395, where we also saw no impact on T6SS sensitivity (Figure 3A).

**Figure 3.**
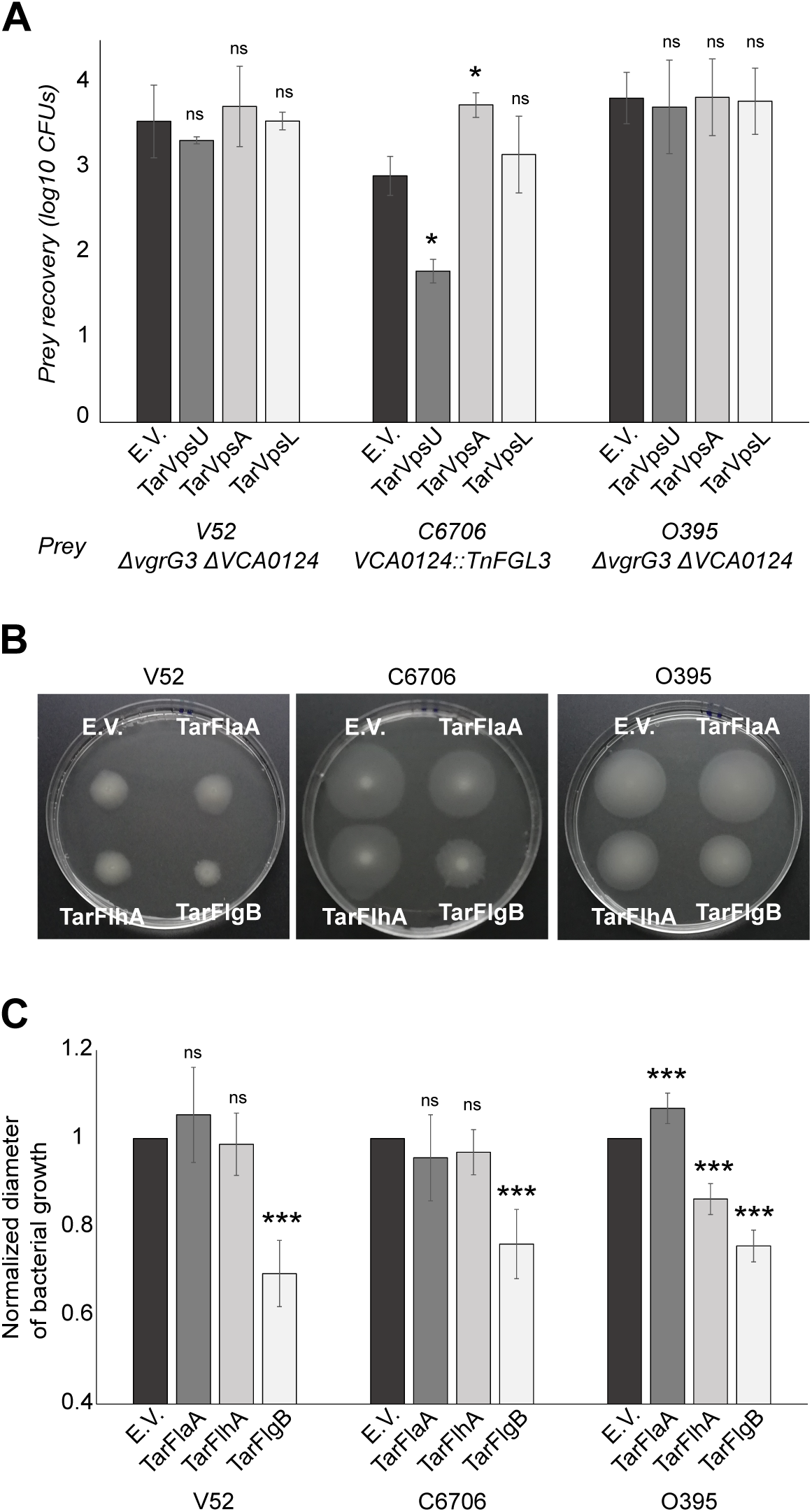
Engineered sRNAs can modulate EPS production and motility in *V. cholerae*. **(A)** Quantification of prey CFUs recovered after the competition assay between *different V. cholerae* strains carrying pBAD33 with sRNAs targeting EPS production (preys) and the *V. cholerae* 2740-80 *clpV*::*clpV*-*mCherry* (predator T6SS +). Data represent the average of at least three independent replicates, each one done in technical duplicates. Error bars represent the SD. E.V., empty vector. **(B)** Representative image of the motility assay of different *V. cholerae* strains carrying sRNAs targeting the flagellum apparatus expressed from pBAD33 backbones. **(C)** Quantification of the motility halos (diameter) shown in panel C. Data is represented as a ratio between the halo of a sample in relation to the halo diameter of the empty vector sample (normalized diameter). Data represent the average of at least four independent replicates, each one done in technical duplicates. Error bars represent the SD of these replicates. Asterisks represent statistical significance when compared to the empty vector sample (paired two-tail t-test, * p value <0.05, ** p value <0.01, ***p value <0.001; ns not significant).

Like biofilm formation, motility is another cellular process that plays a role during both V*. cholerae* infection and environmental growth (35). *V. cholerae* has a single sheathed flagellum encoded by multiple genes transcribed in a hierarchal way and located in three large clusters. Motor genes of the flagellum are the exception as they are in three additional loci (36, 37). We designed three sRNAs targeting different flagellar genes: *flaA* (TarFlaA), *flhA* (TarFlhA) and *flgB* (TarFlgB) (Table S3, File S1). FlaA and FlgB are structural components of the filament and basal body, respectively, while FlhA is involved in the export of the flagellar proteins. *flhA* and *flgB* are the first genes of their flagellar operons, while *flaA* is transcribed as a single transcriptional unit (36, 37). We introduced each of the sRNAs into V52, C6706, and O395 and measured their motility on using a tryptone soft agar plate assay (Figure 3B, C). Strains expressing TarFlgB exhibited less motility relative to the wild type in all three strains, while TarFlhA only affected motility in O395. TarFlaA expression had no effect on V52 or C6706, but slightly increased motility in O395.

Altogether, these results indicate that the efficacy of targeting each specific cellular process will depend on the *V. cholerae* strain. Notably, our engineered sRNAs were most effective in *El tor* O1 strains, where the scaffold TarB sRNA is known to be functional (22, 38).

### Delivery of TarVipA through conjugation results in rapid loss of T6SS activity

Although our engineered sRNAs could successfully knockdown different cellular processes, it was after expression of the sRNA across multiple generations. We next looked to see how quickly after conjugative delivery of an sRNA gene into a given cell would its effects be visible. Since TarVipA had the clearest phenotypic effect among our engineered sRNAs, we focused on this sRNA.

We first cloned the sRNA gene into a mobilizable plasmid carrying the oriT region from the IncP plasmid RP4. To track successful conjugative delivery, we also cloned into the plasmid an array of approximately 100 copies of the Tet operator (*tetO)* (39). The cognate repressor TetR, fused to mNeonGreen, was then constitutively expressed in the recipient *V. cholerae* cells. Upon successful delivery of the sRNA plasmid, the TetR-mNeonGreen present in the cell cytosol would bind to the *tetO* array coalescing into a green fluorescent focus (Figure 4A). For the *V. cholerae* strain, we used the *2740-80 clpV::clpV-mCherry* strain described earlier. When the plasmid carrying the TarVipA and *tetO* array was introduced in *V. cholerae*, although there were still some T6SS activity, there was substantially less activity compared to the repressed control (cells grown in glucose) or a non-targeted control sRNA (TarVCA0688) (Figure S3).

**Figure 4.**
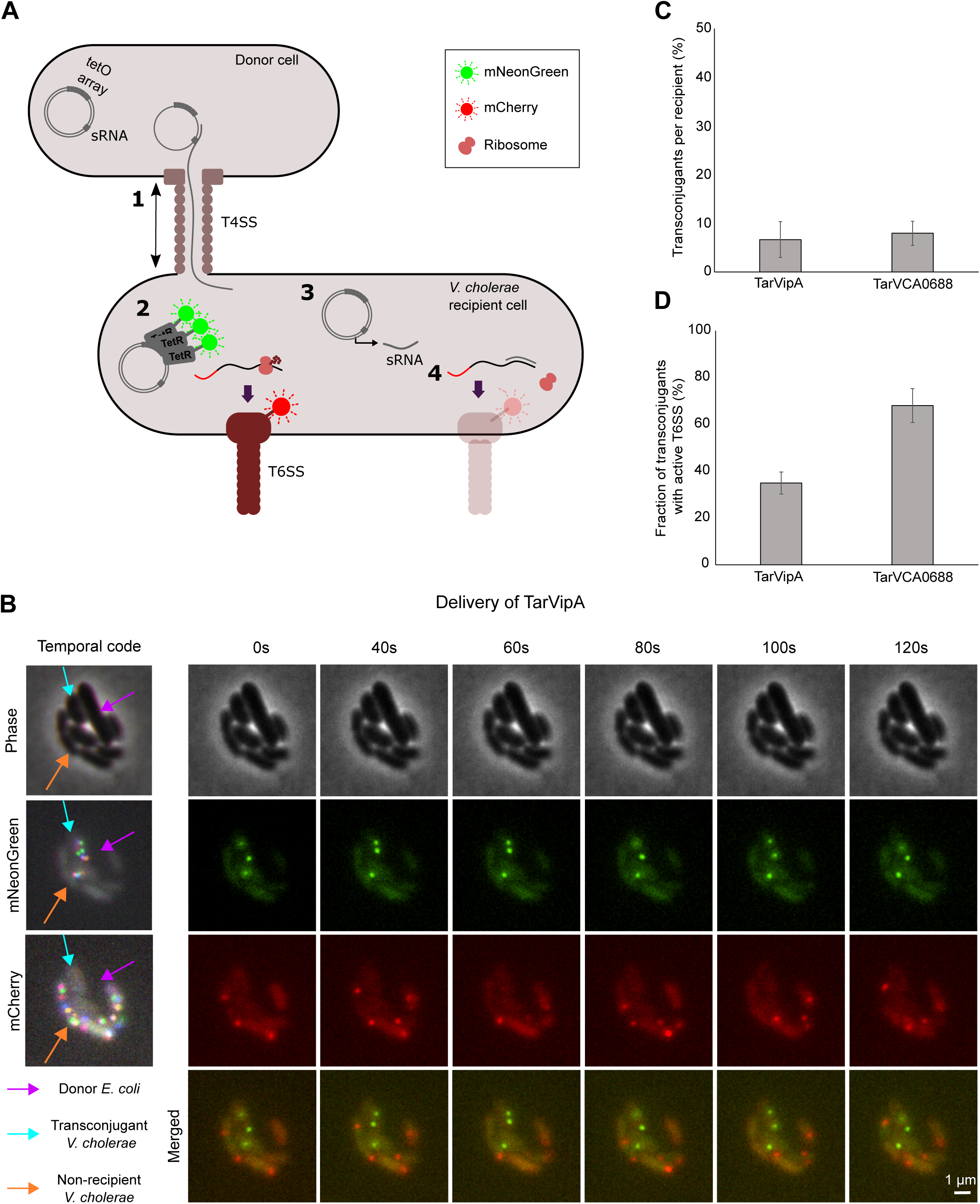
TarVipA rapidly inhibits *V. cholerae* T6SS activity when delivery via conjugation. **(A)** Illustrative image of the assay developed to track conjugation events and T6SS activity with fluorescence microscopy. *E. coli* MFD*pir* donor cells, carrying either pBAD33-TarVipA-Lau44T or pBAD33-TarVCA0688-Lau44T, were mixed with *V. cholerae* 2740-80 *clpV::clpV-mCherry* pBAD33-J23103-tetR-mNeonGreen recipient cells 10:1 and incubated for 90 min before imagining . First, donors transfer the plasmid carrying the *tetO* array and sRNA (TarVipA or TarVCA0688) into a *V. cholerae* strain through the type IV secretion system (T4SS) (1). This *vibrio* strain has a ClpV-mCherry fusion to detect T6SS activity. It also expresses a TetR-mNeonGreen fusion that recognises and binds to the *tetO* array, shifting the green fluorescence from being ubiquitous in the cell to localized in a specific region (focus) (2). Once the plasmid is in the recipient cell it transcribes the sRNA (3) that will bind to its target mRNA, inhibiting its translation. The sRNA function will be observed by a reduction in the T6SS activity (4). Three biological replicates were conducted for each sRNA delivered. **(B)** Representative image of a 2 min time-lapse microscopy showing the delivery of TarVipA sRNA. An *E. coli* donor can be found surrounded by several *V. cholerae* cells, 4 of them in direct contact with the *E. coli* cell. Of those 4, 2 were transconjugants (presence of green foci) The red channel shows the presence of active T6SS foci. Merge images are the combination of the red and green channels. Temporal code image analysis of phase, red and green channels was performed to detect dynamic foci. **(C)** Quantification of transconjugants in relation to the total number of recipient cells for both sRNAs delivered. **(D)** Quantification of conjugated cells with an active T6SS after delivery of TarVipA or TarVCA0688. The difference between these samples was statistically significant (paired two-tail t-test, p value <0.01).

To measure the effect of the conjugatively delivered TarVipA on T6SS activity, we mixed a conjugative donor strain of *E. coli* carrying the sRNA plasmid with recipient *V. cholerae* on a microscope slide, then looked for the presence of dynamic clpV-mCherry foci in transconjugant cells containing TetR-mNeonGreen foci (Figure 4B). We allowed 90 min of incubation prior to microscopy visualization. By that time, 6-7% of all *V. cholerae* cells had acquired the sRNA plasmid (Figure 4C). Notably, the specific sRNA encoded in the plasmid did not affect the rate of conjugation. Of the transconjugants receiving the TarVipA plasmid, only 34% of the cells still had an active T6SSs (Figure 4D). By contrast, when control sRNA TarVCA0688 was delivered roughly 70% of transconjugant cells had an active T6SS. This ∼40% decrease in T6SS activity after only 90 minutes shows that conjugative delivery of sRNA can rapidly modulate cellular process in recipient cells.

## DISCUSSION

Overall, this work represents a proof of concept for the use of conjugatively delivered sRNAs to modulate gene expression in *V. cholerae.* We have engineered sRNAs using TarB from *V. cholerae* as a scaffold to modulate T6SS activity, EPS production, and motility. Notably, disruption of these cellular processes required only the expression of the sRNA without the need for delivery and expression of accessory factors, as is needed for other gene silencing strategies previously used in *V. cholerae* (40). The efficiency of gene expression modulation by these sRNAs ranged from completely inhibiting a cellular process (TarVipA) to having nearly no effect (TarVpsL). However, when knockdown of a given process was successful, we observed a large tolerance for variability of the target recognition site. Unless there was an extreme disruption of the original secondary structure, the gene knockdown was equally effective. Additionally, we found no obvious indicators for how effective a given sRNA would be based on the primary sequence or predicted secondary structure. As such, the variability in the effectiveness of our various sRNAs is likely intrinsic to the target genes themselves. This conclusion is further supported by our observation that the same sRNA could have substantially different efficacy in different *V. cholerae* strains. For example, an sRNA targeting exopolysaccharide production (TarVpsU) was highly effective in El Tor O1 strain, C6706, but the same sRNA had no effect in the Classical O1 strain, O395. Similarly, an sRNA targeting motility (TarFlhA) was effective in O395 but not in C6706. These strain-specific differences are probably due to differences in the way these systems are regulated in each strain. Therefore, to ensure effective knockdown of a given cellular system, future more practical applications may require concomitant delivery of multiple sRNAs targeting different genes within a given system. That said, such strain specific effects may actually be advantageous in situations where only a subset of strains in a multi-strain community needs to be targeted.

One advantage of using sRNAs to modulate genes (as opposed to a system like dCas9) is that because it exploits an existing regulatory system, only the delivery and expression of the sRNA itself is required. As such, targeted bacteria do not need to be pre-loaded with accessory regulatory factors, allowing knockdown effects to occur at the rate of protein turnover. In our work, we were able to gain insight into the approximate kinetics of target system modulation by simultaneously tracking conjugative delivery of the sRNA gene and the subsequent shutdown of T6SS activity. We were able to observe sRNA knockdown effects occurring within a single cell division cycle following the conjugation event. Given such rapid response rates, the limiting factor for microbial community modulation will be the rate of sRNA delivery.

The broad host-range conjugative systems employed in this study are an attractive option because they enable cross-species delivery from commensal bacteria into *V. cholerae*. Indeed, contact-dependent antagonistic interactions of *V. cholerae* with resident commensal bacteria mediated by the T6SS have been shown to play a role in *V. cholerae* pathogenesis (41). Given that *V. cholerae* is making direct physical contact with these bacteria, pre-loading them with a conjugative element carrying an anti-T6SS sRNA could be an effective delivery strategy so long as the conjugative element could be stably maintained within the resident population. However, the conjugation rates we observed with our broad host-range conjugative system are probably too low to enable their use for delivery of regulatory sRNAs into larger established communities.

While, more efficient delivery may be possible with a Vibrio-specific conjugative plasmid like P factor (14), temperate bacteriophages have already been demonstrated to be an effective delivery tool for *in situ* modification of microbial communities (11, 12). However, the high specificity afforded by phage-based delivery systems necessarily means a narrow range of targetable strains, limiting their use prophylactically. On the other hand, broad-host range conjugative elements can be delivered into a wide range of bacterial species shifting the selectivity and specificity to the sRNA gene and its regulation. As such, incorporating conjugative elements carrying engineered sRNA genes into either probiotic or live vaccine strains should be feasible, but further studies using *in vivo* infection models will be needed to fully assess their efficacy.

## AUTHOR CONTRIBUTIONS

P.M.-G. and B.T.H conceived and designed the experiments. P.M.-G., D.V. and R.R. performed the experiments. P.M.-G., and B.T.H analysed the data and wrote the manuscript. B.T.H acquired funding for this project.

## ACKNOWLEDGEMENT

This work was supported by MRC grant MR/T031131/1.

